# Quantifying orthogonal barcodes for sequence census assays

**DOI:** 10.1101/2022.10.09.511501

**Authors:** A. Sina Booeshaghi, Kyung Hoi (Joseph) Min, Jase Gehring, Lior Pachter

## Abstract

Barcode-based sequence census assays utilize custom or random oligonucloetide sequences to label various biological features, such as cell-surface proteins or CRISPR perturbations. These assays all rely on barcode quantification, a task that is complicated by barcode design and technical noise. We introduce a modular approach to quantifying barcodes that achieves speed and memory improvements over existing tools. We also introduce a set of quality control metrics, and accompanying tool, for validating barcode designs.

## Introduction

Single-cell RNA sequencing assays rely on unique sequences of nucleotides (“barcodes”) to associate sequences from mRNA molecules with individual cells. This strategy has been extended to quantify orthogonal biological or technical information in what is known as “multimodal” genomics. For example, the 10x Genomics Feature Barcoding assay quantifies the expression of cell-surface proteins using targeted antibody barcodes (“Feature Barcode-Overview -Software -Single Cell Gene Expression -Official 10x Genomics Support” n.d.). The ClickTag (Gehring et al. 2020), Multiseq (McGinnis et al. 2019), and Cell Hashing (Stoeckius et al. 2018) assays, on the other hand, use custom sample-specific barcodes to group cells enabling multiplexing and reducing batch effects. Moreover, similar approaches have been used for CRISPR screens (Dixit et al. 2016; Datlinger et al. 2017; Schraivogel et al. 2020), targeted perturbation assays (Gehring et al. 2020; Srivatsan et al. 2020), and recently for spatial genomics assays (Srivatsan et al. 2021).

The processing of data from these assays has required the development of custom tools, resulting in a proliferation of application- and assay-specific software (Roelli et al. 2019; Milo S. Johnson, Sandeep Venkataram, Sergey Kryazhimskiy 2022). We introduce a broadly applicable framework, which we term *“kITE”* for ***k**allisto* **I**ndexing and **T**ag **E**xtraction, for uniformly quantifying orthogonal barcode-based sequence census assays that has several advantages over assay-specific tools. First, by virtue of providing a single solution to seemingly distinct problems, it serves as a unifying framework for *biology x barcode* assays and will simplify the development and deployment of novel assays. Second, the software implementation we share is portable, engineered to work across platforms, and is modular, allowing for easy adaptation and customization. It makes use of the *kallisto* and *bustools* programs that have been previously extensively validated and benchmarked for quantification of the biological sequences in barcode x biology assays such as single-cell and bulk RNA sequencing (Bray et al. 2016; Melsted et al.2021). Finally, our method is faster than existing tools, and we identify corner cases in processing that have been missed in previous analyses. The examination of such cases led us to develop a tool, called *qcbc,* for assessing the quality of barcode sets, and that should prove to be a useful standalone package. It is applicable both during assay development, and during assay quality control.

### *kITE* design

Feature Barcoding assays use short barcodes and are susceptible to ambiguous or incorrect assignment if errors are introduced during the experimental procedure. Inspired by the *CITE-seq-count* approach, *kITE* begins by generating a list of all single-base mismatches of the Feature Barcodes used in an experiment (**Figure 1**). The resulting ‘mismatched FASTA’ file is used as input for the *‘kallisto* index’ program with *k* set appropriately (methods). Finally, the *kallisto* | *bustools* pipeline is used to quantify the dataset using the ‘mismatch index’ generated after running *‘kallisto* index’ on the mismatched FASTA file. In this way, *kallisto* will effectively scan the entire sequencing read for barcodes present in the mismatch index allowing for barcode pseudoalignment despite variability in barcode position. Feature barcode “error correction” occurs when barcode counts are summed across different mismatches of a given barcode (Methods).

**Figure 1:**
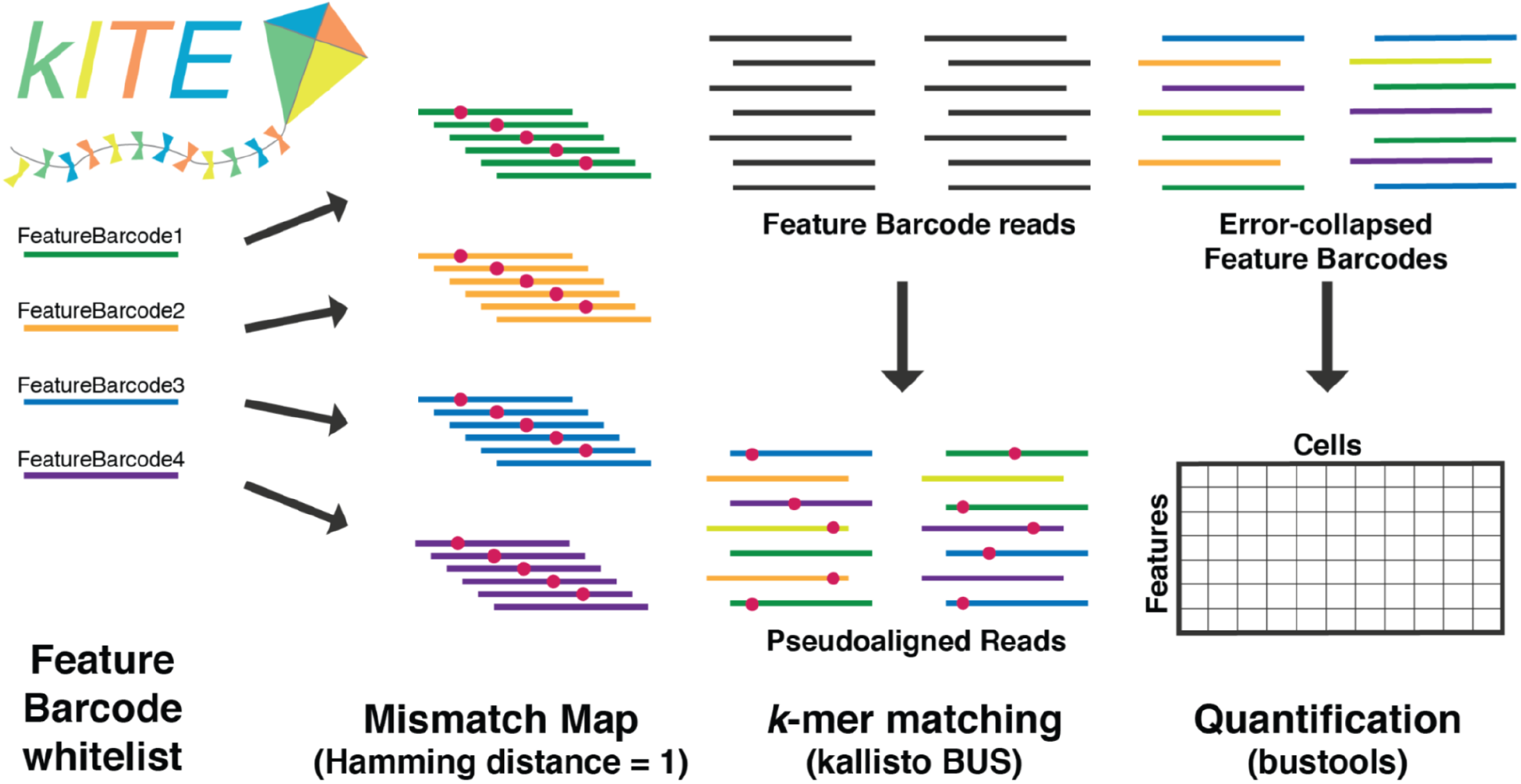
*kITE* workflow. The mismatch map is generated from the feature barcode whitelist and indexed with *kallisto.* Reads are pseudoaligned and error-collapsed with *kallisto* and *bustools* to form a cells x features matrix.

### Method comparison

To evaluate the accuracy and efficiency of *kITE* across a range of assays, we gathered data from six different assays, and compared *kITE* quantifications to those produced with assay-specific tools. We first quantified cell surface protein abundance for the 10x Genomics 5,266 Peripheral blood mononuclear cells dataset, which targeted 32 cell surface proteins. We used *kallisto bustools* and a mismatch index to pseudoalign the target sequences (**Methods: Index, pseudoalignment, and counting**) and found high concordance between our method and *Cell Ranger* across varying read depths, with an average Pearson correlation of 0.99 (**Figure 2a-f**). *kiTE* was five times faster than *Cell Ranger* and required 3.5 times less RAM (**Methods: Speed and memory comparison**).

**Figure 2:**
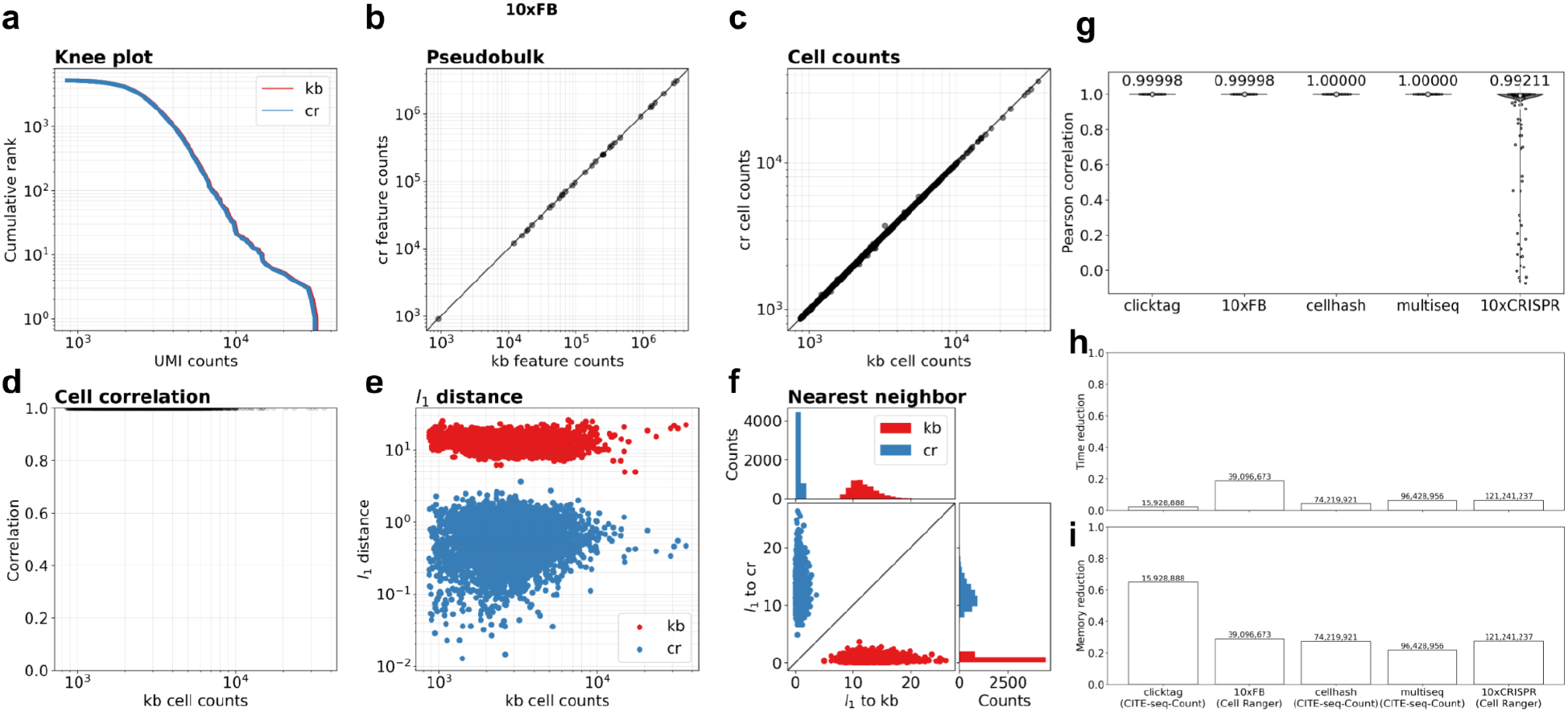
Comparison between *kallisto bustools* (kb) quantifications and Cell Ranger (cr) quantifications for the 10x Feature Barcoding assay on 5,266 Peripheral blood mononuclear cells dataset targeting 32 cell surface proteins. **(a)** Knee plot comparing cumulative UMI counts per cell. **(b)** Pseudobulk comparison of cumulative UMI counts per feature barcode. **(c)** Cumulative UMI counts per cell. **(d)** Pearson correlation of the same cell between the two quantifications. **(e)** The *l*_1_ distance between a *kallisto bustools* cell and its Cell Ranger equivalent (blue) and the same cell and its nearest *kallisto bustools* neighbor (red) across the total UMI counts for that cell. **(f)** The *l*_1_ distance of a *kallisto bustools* cell (red) to its Cell Ranger equivalent (y-axis) and to its nearest neighbor (x-axis) and the *l*_1_ distance of a Cell Ranger cell (blue) to it’s *kallisto bustools* equivalent (x-axis) and its nearest neighbor (y-axis). The marginal distributions show that each *kallisto bustools* cell is closest to its corresponding Cell Ranger cell and that each Cell Ranger cell is nearest to its corresponding *kallisto bustools* cell. **(g)** Pearson correlation for all the cells between the *kallisto bustools* qualifications and in *CITE-seq-Count* or *Cell Ranger* quantifications for each of the five datasets. **(h)** Runtime and **(i)** memory improvements for each of the five datasets across the various read depths.

Next, we tested how accurately we could quantify and demultiplex pooled samples tagged with sample-specific barcodes. We processed data from 3,795 methanol-fixed mouse neural stem cells from four multiplexed samples generated with the ClickTag assay (**Supplementary figure 1**), 9,551 primary patient-derived xenograft (PDX) samples from nine multiplexed samples generated with the Multiseq assay (**Supplementary Figure 2)**and 10,807 PBMCs from eight multiplexed samples generated with the Cell Hashing assay (**Supplementary Figure 3**). We found high concordance with an average Pearson correlation of 0.99, for all three datasets, between *kITE* and *CITE-seq-Count* (Roelli et al. 2019), a tool for counting antibody tags from a CiteSeq or Cell Hashing experiments (**Figure 2g**). *kITE* was 48 times faster and required only a quarter to a fifth of the RAM (**Figure 2h,i**). We observed slight discordance with two feature barcodes, BC49 and BC74, when processing the PDX samples assayed with Multiseq. These two Multiseq barcodes are short (8 bp in length), yet share long subsequences up to 7 bp in length **(Supplementary Figure 4a)**. The introduction of errors during sequencing can result in possible incorrect barcode assignment and loss of barcodes (**Methods: Barcode simulation**). We addressed this issue by trimming the excess sequence surrounding the 8 bp feature barcode (**Methods: Multiseq**).

To demonstrate the flexibility of our approach we also processed 2,799 A549 lung carcinoma cells expressing nuclease-dead Cas9–Krüppel-associated box (dCas9–KRAB) assayed with the 10x Genomics CRISPR screen kit, and transduced with 90 sgRNAs targeting 45 different genes. We achieve high concordance against *Cell Ranger* with an average Pearson correlation of 0.99 across all cells for nearly sgRNA’s, with two sgRNAs in the 10x CRISPR screen dataset, EZR-1 and PPIB-2, differing substantially (**Supplementary Figure 5**). Both of these two sgRNA’s share a subsequence of 16 bp with EZR-2 and PPIB-1 respectively. The latter two sequences begin with GCG and GGA, both of which are a single hamming distance away from the three bases that precede the target sgRNA in the protospacer sequence (**Supplementary Figure 4b**). Therefore a single mutation two bases away from the start of the EZR-1 sgRNA sequence could result in an ambigous read. To resolve this we trimmed the protospacer sequence off of the reads (**Methods: 10xCRISPR**).

We also processed 10,860 K562 cells expressing dCas9–KRAB from TAP-Seq transduced with 86 sgRNAs targeting 14 promoters or enhancers and 30 control sgRNAs to compare assignment of sgRNAs to individual cells. We found 97.8% of all 10,860 barcodes are fully concordant with TAP-Seq assignments, 0.53% of cells are partially concordant, and 1.6% of cells are fully discordant. (**Methods: TAP-Seq**). The discrepancies are likely the result of differences between mapping procedures producing differences in counts near the threshold (8 umi counts) resulting in differential cell assignment for cells with low UMI counts (**Supplementary Figure 6)**.

### Read trimming

Sequencing libraries often contain adapter or spurious sequences that flank the barcode and the presence of these shared sequences can make mapping challenging. Sequencing reads are often trimmed, with tools such as trimmomatic (Bolger, Lohse, and Usadel 2014), to avoid mapping non-informative regions that flank a barcode, a task that is crucial when barcodes share subsequences at their ends. Trimming helps to avoid longer sequence clashes that contain both the shared barcode subsequence and the shared non-barcode sequence. The 10xCRISPR assay, for example, contains two barcodes which share a 16bp subsequence (the ends of which are one hamming distance from a subsequence of the protospacer sequence that flask the barcode (**Supplementary Figure 4**). Similarly, the Multiseq assay produces sequencing reads where the barcodes are flanked by poly-A stretches and two barcodes share a 7bp subsequence that contains the end of the barcode.

### Barcode design validation

Orthogonal barcode sequences designed for quantifying multimodal data, demultiplexing samples, or performing large-scale perturbation experiments can clash if the sequences are not sufficiently unique. To allow researchers to assess the extent of these issues in datasets they are analyzing, or for assays they are developing, we developed a quality control tool, *qcbc,* and a simulation framework to assess the quality of a set of barcodes that is complementary to the simulation framework proposed in (Milo S. Johnson, Sandeep Venkataram, Sergey Kryazhimskiy 2022). *qcbc* computes multiple metrics for barcode quality control:

1. diversity of unique barcodes (**Figure 3a**),
2. number of barcodes that share a subsequence of a given length (**Figure 3b**),
3. distribution of Hamming distances between pairwise barcodes (**Figure 3c,d**),
4. distribution of homopolymers of a given length (**Figure 3e**), and
5. per position and barcode nucleotide distribution (**Figure 3f, Supplementary Figure 7**).

**Figure 3:**
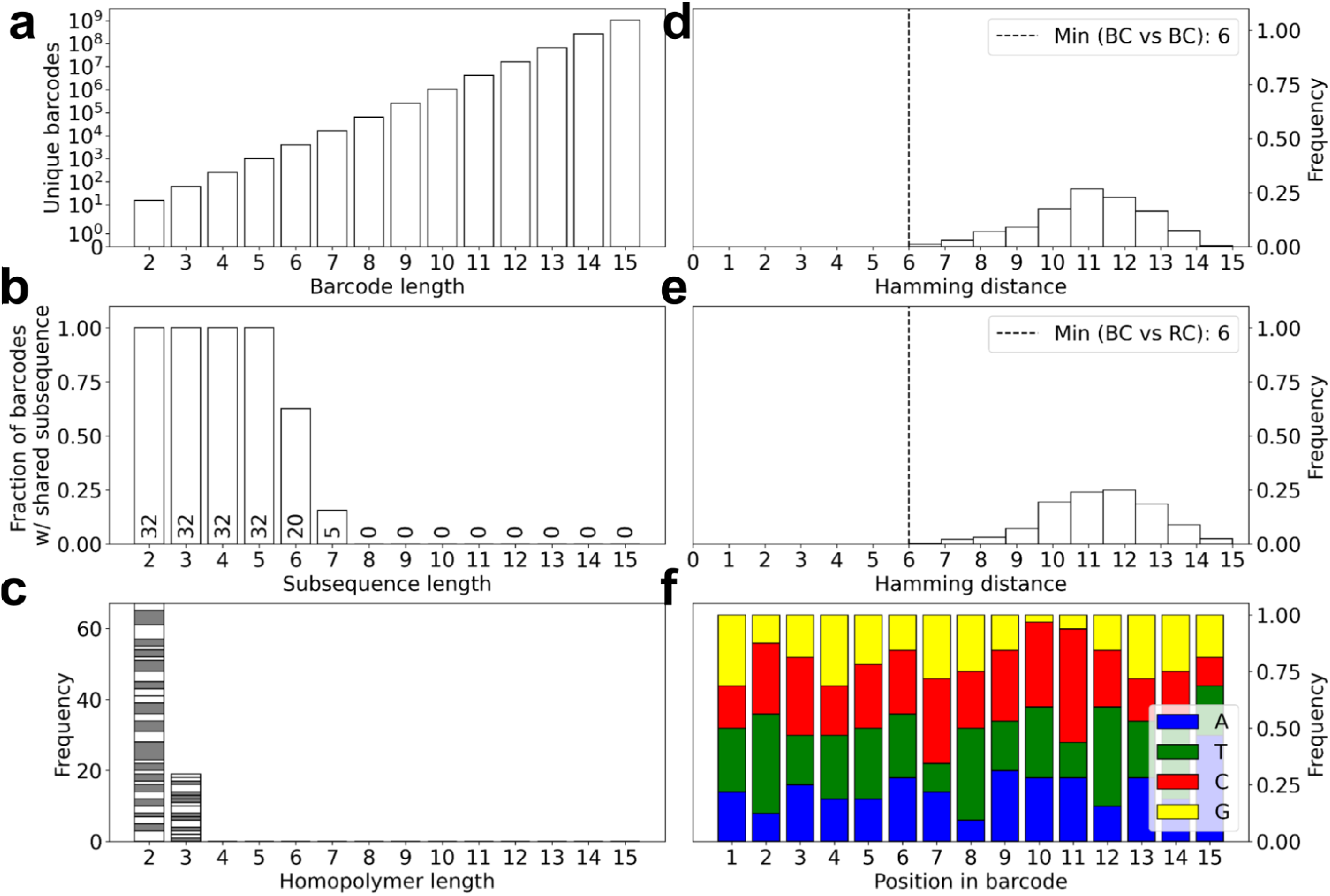
Evaluation of barcode designs. **(a)** Potential library diversity for 4-letter DNA barcodes of various lengths. **(b)** Number of ambiguous barcodes for varying subsequence length. **(c)** The distribution of the number of homopolymer stretches of a given length. The different color bands indicate a unique barcode and the size of the band indicates the number of homopolymers of that length found in that barcode. **(d)** Distribution of pairwise hamming distances between barcodes. The minimum pairwise Hamming distance is marked with a dashed line. **(e)** Distribution of pairwise hamming distances between barcodes and their reverse complement. The minimum pairwise Hamming distance is marked with a dashed line. **(f)** The per base nucleotide content across all barcodes.

With *qcbc* we are able to identify the 16 bp shared subsequence with EZR-2 and PPIB-1 sgRNAs in the 10xCRISPR that result in improper quantification. *qcbc* is also able to identify the 7bp shared subsequence between BC49 and BC74 in the Multiseq assay.

These metrics are important for understanding the impact that barcode error correction has on rescuing sequencing reads. A large diversity of unique barcodes with shorter shared subsequences and larger pairwise distances between barcodes make it easier to disambiguate sequencing reads with pseudoalignment. Additionally, long homopolymer runs and low nucleotide content diversity could contribute to sequencing errors (Sina Booeshaghi and Pachter 2022; “What Is Nucleotide Diversity and Why Is It Important?” n.d.).

To evaluate the impact that base errors have on error correction, we developed a simulation framework that reports the fraction of lost barcodes for a given error rate in the observed sequences **Supplementary Figure 8** (**Methods: Barcode simulation**). Short barcodes, such as those in the Multiseq assay can yield insufficient barcode diversity that results in a greater fraction of barcodes that cannot be unambiguously assigned **Supplementary Figure 9**. Barcodes with small minimum-pairwise / edit distance **Figure 3d,e** and long shared subsequences **Figure 3b** such as those in the Mulitseq and 10xCRISPR assay can result in more clashes (**Methods: Barcode validation**).

## Discussion

We have demonstrated a broadly applicable approach to align, error-correct, group, and count sequences from orthogonal barcodes to quantify multimodal data such as cell surface protein abundance, to demultiplex samples, and to perform large-scale perturbation experiments. Our approach is fast, memory efficient, and extensible: orthogonal barcode quantifications can be obtained for numerous assays with variable sequence structure such the recently developed spatial RNA-seq assays (Srivatsan et al. 2021) and massively parallel reporter assays (Gordon et al. 2020). This allows future assay developers to avoid producing, validating, and maintaining software implementing *ad hoc* solutions. Moreover, some current methods are slow, have high memory requirements, and are specifically tailored for specific assays, which can result in batch effects when performing integrative analysis. Methods such as *CITE-seq-Count* offer extensibility features for different barcode strategies, but are slow and memory-intensive. The speed and efficiency of our approach enables reproducible single-cell analysis in the cloud with tools such as Google Colab. We anticipate that our framework for quantifying orthogonal barcoding assays will be useful as multimodal assays proliferate (Packer and Trapnell 2018), and that our quality control tool will help assay developers avoid pitfalls in barcode design and help practitioners quality control orthogonal barcode assays. Finally, while our approach is currently engineered to correct mismatch errors, robustness to other types of sequencing errors are possible via other methods that could readily be incorporated in our modular framework (Zorita, Cuscó, and Filion 2015; Booeshaghi et al. 2020; Milo S. Johnson, Sandeep Venkataram, Sergey Kryazhimskiy 2022).

## Methods

All code, parameter options, and data can be found in the paper’s GitHub repository: https://github.com/pachterlab/BMGP_2020. *kite* is implemented in *kb_python* and can be found here: https://github.com/pachterlab/kb_python. The *qcbc* barcode validator can be found: https://github.com/pachterlab/qcbc.

### Mismatch index, pseudoalignment, and counting

To quantify against a known set of target sequences we first build an alignment index containing all Hamming distance = 1 variants of each target. Restricting the alignment index to Hamming distance = 1 variants allows for recovering more reads while keeping the index size small **Supplementary Figure 10**. The hamming-1 index for the Clicktag assay is 72.8 KB and the hamming-0 index is 2.1 KB. Targeted sequencing reads are then pseudoaligned to the custom index using *kallisto* to generate a Barcode, UMI, Set (BUS) file(Melsted, Ntranos, and Pachter 2019). UMI counts are then aggregated per cell and quantified per feature barcode using *bustools* to generate a target count matrix.

### Speed and memory comparison

Preprocessing steps were benchmarked with GNU time -v option, and the elapsed (wall clock) time and maximum resident set size (kbytes) were used to perform the time and memory comparisons. All code was benchmarked on Ubuntu 18.04.5 LTS (GNU/Linux 4.15.0-136-generic x86_64) with Intel(R) Xeon(R) CPU E5-2697 v2 @ 2.70GHz processors.

### 10x Feature Barcode

FASTQ files for 5,266 Peripheral blood mononuclear cells dataset targeting 32 cell surface proteins were downloaded from https://support.10xgenomics.com/single-cell-gene-expression/datasets/3.0.2/5k_pbmc_protein_v3. A mismatch index was created using the Feature Reference file and *kb* ref with the *kite* workflow. The reads were pseudoaligned and a feature count matrix was made using *kb* count with the kite:10xFB technology option. *Cell Ranger,* the Feature Reference, and refdata-gex-GRCh38-2020-A was used to process the FASTQ files into a feature count matrix.

All preprocessing steps are runnable via a Google Colab notebook: https://github.com/pachterlab/BMGP_2020/blob/main/analysis/notebooks/10xFB/10xFB_preprocess.ipynb. All matrix comparisons are runnable via a Google Colab notebook: https://github.com/pachterlab/BMGP_2020/blob/main/analysis/notebooks/10xFB/10xFB_preprocess.ipynb

### ClickTag

FASTQ files for 3,795 methanol-fixed mouse neural stem cells from four multiplexed samples were downloaded from https://caltech.box.com/shared/static/zqaom7yuul7ujetqyhnd4lvf8vhzqsyg.gz. A mismatch index was created using the list of targeted barcodes file and *kb* ref with the *kite* workflow. The list of barcodes can be found here: https://github.com/pachterlab/BMGP_2020/tree/main/references/clicktag. The reads were pseudoaligned and a feature count matrix was obtained using *kb* count with the *kite* technology option. *CITE-seq-Count* and the targeted barcodes were used to process the FASTQ files into a feature count matrix.

All preprocessing steps are runnable via a Google Colab notebook: https://github.com/pachterlab/BMGP_2020/blob/main/analysis/notebooks/clicktag/clicktag_preprocess.ipynb. All matrix comparisons are runnable via a Google Colab notebook: https://github.com/sbooeshaghi/BMGP_2020/blob/main/analysis/notebooks/clicktag/clicktag_preprocess.ipynb

We generated a *kallisto* index with no mismatches by running *kb* ref with the --no-mismatches option and compared the resultant count matrix generated with *kb* count to the count matrix generated with mismatches.

### Multiseq

FASTQ files for 9,551 primary patient-derived xenograft (PDX) samples from nine multiplexed samples were downloaded from https://caltech.box.com/shared/static/0scoe38xvcoxfd62xnm848pimh2zr1ip.gz. A mismatch index was created using the list of targeted barcodes file and *kb* ref with the *kite* workflow. The list of barcodes can be found here: https://github.com/pachterlab/BMGP_2020/tree/main/references/multiseq. The reads were pseudoaligned and a feature count matrix was obtained using *kb* count with the *kite* technology option. *CITE-seq-Count* and the targeted barcodes were used to process the FASTQ files into a feature count matrix.

All preprocessing steps are runnable via a Google Colab notebook: https://github.com/pachterlab/BMGP_2020/blob/main/analysis/notebooks/multiseq/multiseq_preprocess.ipynb. All matrix comparisons are runnable via a Google Colab notebook: https://github.com/sbooeshaghi/BMGP_2020/blob/main/analysis/notebooks/multiseq/multiseq.ipynb

The FASTQ reads were trimmed using *seqtk* trimfq with the -e option set to the length of the read minus the 8 such that only the 8 bp barcode was retained.

### Cell Hashing

FASTQ files for 10,807 PBMCs from eight multiplexed samples were downloaded from https://caltech.box.com/shared/static/0scoe38xvcoxfd62xnm848Dimh2zr1ip.qz. A mismatch index was created using the list of targeted barcodes file and *kb* ref with the *kite* workflow. The list of barcodes can be found here: https://github.com/pachterlab/BMGP_2020/tree/main/references/multiseq. The reads were pseudoaligned and a feature count matrix was obtained using *kb* count with the *kite* technology option. *CITE-seq-Count* and the targeted barcodes were used to process the FASTQ files into a feature count matrix.

All preprocessing steps are runnable via a Google Colab notebook: https://github.com/pachterlab/BMGP_2020/blob/main/analysis/notebooks/cellhash/cellhash_preprocess.ipynb. All matrix comparisons are runnable via a Google Colab notebook: https://github.com/pachterlab/BMGP_2020/blob/main/analysis/notebooks/cellhash/cellhash.ipynb

### 10x CRISPR

FASTQ files for 2,799 A549 lung carcinoma cells expressing nuclease-dead Cas9–Krüppel-associated box (dCas9–KRAB) from 10x Genomics CRISPR screen transduced with 90 sgRNAs targeting 45 different genes were downloaded from https://support.10xgenomics.com/single-cell-gene-expression/datasets/4.0.0/SC3_v3_NextGem_DI_CRISPR_10K. A mismatch index was created using the Feature Reference file and *kb* ref with the *kite* workflow. The reads were pseudoaligned and a feature count matrix was obtained using *kb* count with the kite:10xFB technology option. *Cell Ranger,* the Feature Reference, and refdata-gex-GRCh38-2020-A was used to process the FASTQ files into a feature count matrix.

All preprocessing steps are runnable via a Google Colab notebook: https://github.com/pachterlab/BMGP_2020/blob/main/analysis/notebooks/10xCRISPR/10xCRISPR_preprocess.ipynb. All matrix comparisons are runnable via a Google Colab notebook: https://github.com/pachterlab/BMGP_2020/blob/main/analysis/notebooks/10xCRISPR/10xCRISPR.ipynb

The FASTQ reads were trimmed using *seqtk* trimfq with the -e option set to the length of the read minus the 19 plus overhang such that only the 19 bp barcode plus the set overhang was retained.

### TAP-Seq

FASTQ files for 10,860 K562 cells expressing dCas9–KRAB from TAP-Seq transduced with 86 sgRNAs targeting 14 promoters or enhancers and 30 control sgRNAs were downloaded from https://www.ncbi.nlm.nih.gov/sra/?term=SRR9916613. A mismatch index was created using the list of targeted barcodes file and *kb* ref with the *kite* workflow. The list of barcodes can be found here: https://github.com/pachterlab/BMGP_2020/tree/main/references/tapseq. The reads were pseudoaligned and a feature count matrix was obtained using *kb* count with the standard technology option. Perturbation status for each cell was downloaded from https://www.ncbi.nlm.nih.gov/geo/query/acc.cgi?acc_GSM4012688 using *ffq* GSM4012688 (Gálvez-Merchán et al. 2022).

All preprocessing steps are runnable via a Google Colab notebook: https://github.com/pachterlab/BMGP_2020/blob/main/analysis/notebooks/tapseq/tapseq_preprocess.ipynb. All matrix comparisons are runnable via a Google Colab notebook:

https://github.com/pachterlab/BMGP_2020/blob/main/analysis/notebooks/tapseq/tapseq.ipynb

### Barcode validation

For a given list of barcodes we report the maximum number of barcodes that could be made for the given barcode length, the number of barcodes that share a subsequence of length k, and the pairwise hamming distance between all pairs of barcodes. A Google Colab notebook reproducing the results of this simulation can be found here: https://github.com/pachterlab/BMGP_2020/blob/main/analysis/notebooks/barcode_validator.jpynb

The *qcbc* tool for barcode validation can be found here: https://github.com/pachterlab/qcbc

### Barcode simulation

We assumed a per-base error rate that is constant for every base in a barcode and simulated barcode mutants from a set of given barcodes. Then, for multiple error rates, we estimated the number of barcode mutants that are a given hamming distance from the correct barcode that became ambiguous after introducing sequencing errors. A Google Colab notebook reproducing the results of this simulation can be found here: https://github.com/pachterlab/BMGP_2020/blob/main/analysis/notebooks/barcode_sim.ipynb

## Software versions

[Anndata 0.6.22.post1](https://github.com/theislab/anndata)

[bustools 0.41.0](https://github.com/BUStools/bustools/)

[Cell Ranger 6.0.1](https://github.com/10XGenomics/cellranger)

CITE-seq-Count 1.4.4

[awk (GNU awk) 4.1.4](https://www.gnu.org/software/gawk/manual/gawk.html)

[grep (GNU grep) 3.1](https://www.gnu.org/software/grep/manual/grep.html)

[kallisto 0.48.0](https://github.com/pachterlab/kallisto)

kb_python 0.26.3

[Matplotlib 3.0.3](https://github.com/matplotlib/matplotlib)

[Numpy 1.18.1](https://github.com/numpy/numpy)

[Pandas 0.25.3](https://github.com/pandas-dev/pandas)

[Scipy 1.4.1](https://github.com/scipy/scipy)

[sed (GNU sed) 4.4](https://www.gnu.org/software/sed/)

[seqtk 1.2-r94](https://github.com/lh3/seqtk)

[sklearn 0.22.1](https://github.com/scikit-learn/scikit-learn)

[tar (GNU tar) 1.29](https://www.gnu.org/software/tar/)

