## Supplementary Information for "Quantifying orthogonal barcodes for sequence census assays"

### **Supplementary Information:**
### Quantifying orthogonal barcodes for sequence census assays

A. Sina Booeshaghi[1], Kyung Hoi (Joseph) Min[2], Jase Gehring[3] and Lior Pachter[1,4]

1. Division of Biology and Biological Engineering, California Institute of Technology, Pasadena, California
2. Department of Computer Science and Electrical Engineering, Massachusetts Institute of Technology, Cambridge, Massachusetts
3. Arcadia Science, Berkeley, California
4. Department of Computing and Mathematical Sciences, California Institute of Technology, Pasadena, California

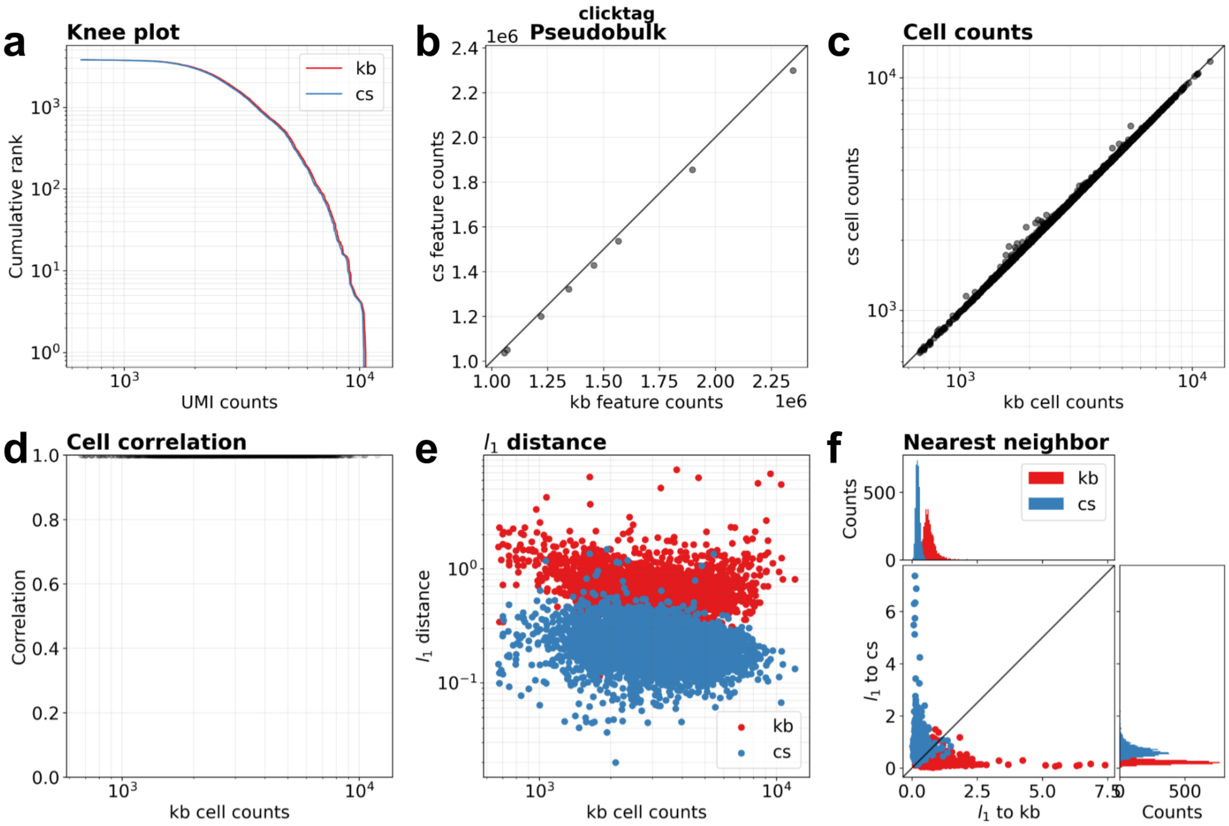


**Supplementary Figure 1:** Comparison between kallisto bustools (kb) quantifications and CITE-seq-Count (cs) quantifications for the 10x Feature Barcoding assay. **(a)** Knee plot comparing cumulative UMI counts per cell. **(b)** Pseudobulk comparison of cumulative UMI counts per feature barcode. **(c)** Cumulative UMI counts per cell. **(d)** Pearson correlation of the same cell between the two quantifications. **(e)** The l_1 distance between a kallisto bustools cell and its CITE-seq-Count doppelganger (blue) and the same cell and its nearest kallisto bustools neighbor (red) across the total UMI counts for that cell. **(f)** The l_1 distance of a kallisto bustools cell (red) to its CITE-seq-Count doppelganger (y-axis) and to its nearest neighbor (x-axis) and the l_1 distance of a CITE-seq-Count cell (blue) to its kallisto bustools doppelganger (x-axis) and its nearest neighbor (y-axis). The marginal distributions show that each kallisto bustools cell is closest to its corresponding CITE-seq-Count cell and that each CITE-seq-Count cell is nearest to its corresponding kallisto bustools cell.

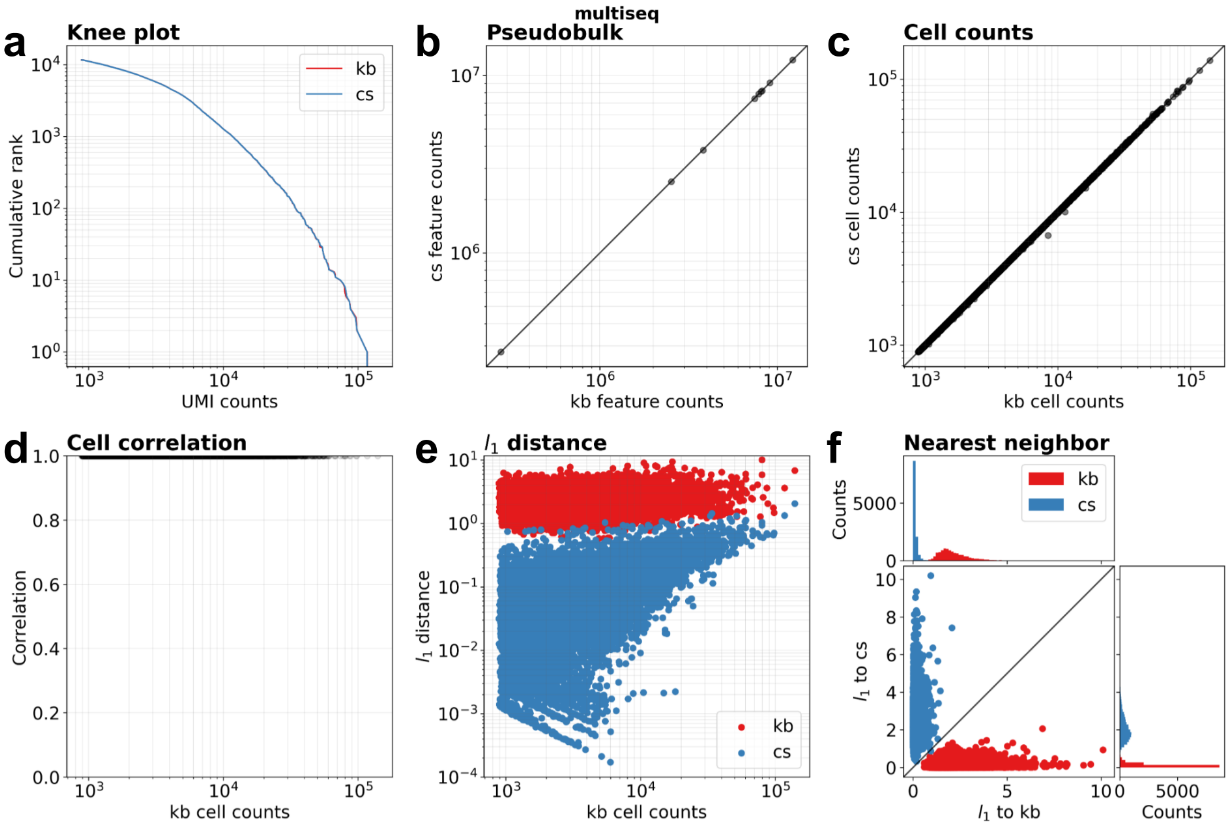


**Supplementary Figure 2:** Comparison between kallisto bustools (kb) quantifications and CITE-seq-Count (cs) quantifications for the 10x Feature Barcoding assay. **(a)** Knee plot comparing cumulative UMI counts per cell. **(b)** Pseudobulk comparison of cumulative UMI counts per feature barcode. **(c)** Cumulative UMI counts per cell. **(d)** Pearson correlation of the same cell between the two quantifications. **(e)** The l_1 distance between a kallisto bustools cell and its CITE-seq-Count doppelganger (blue) and the same cell and its nearest kallisto bustools neighbor (red) across the total UMI counts for that cell. **(f)** The l_1 distance of a kallisto bustools cell (red) to its CITE-seq-Count doppelganger (y-axis) and to its nearest neighbor (x-axis) and the l_1 distance of a CITE-seq-Count cell (blue) to its kallisto bustools doppelganger (x-axis) and its nearest neighbor (y-axis). The marginal distributions show that each kallisto bustools cell is closest to its corresponding CITE-seq-Count cell and that each CITE-seq-Count cell is nearest to its corresponding kallisto bustools cell.

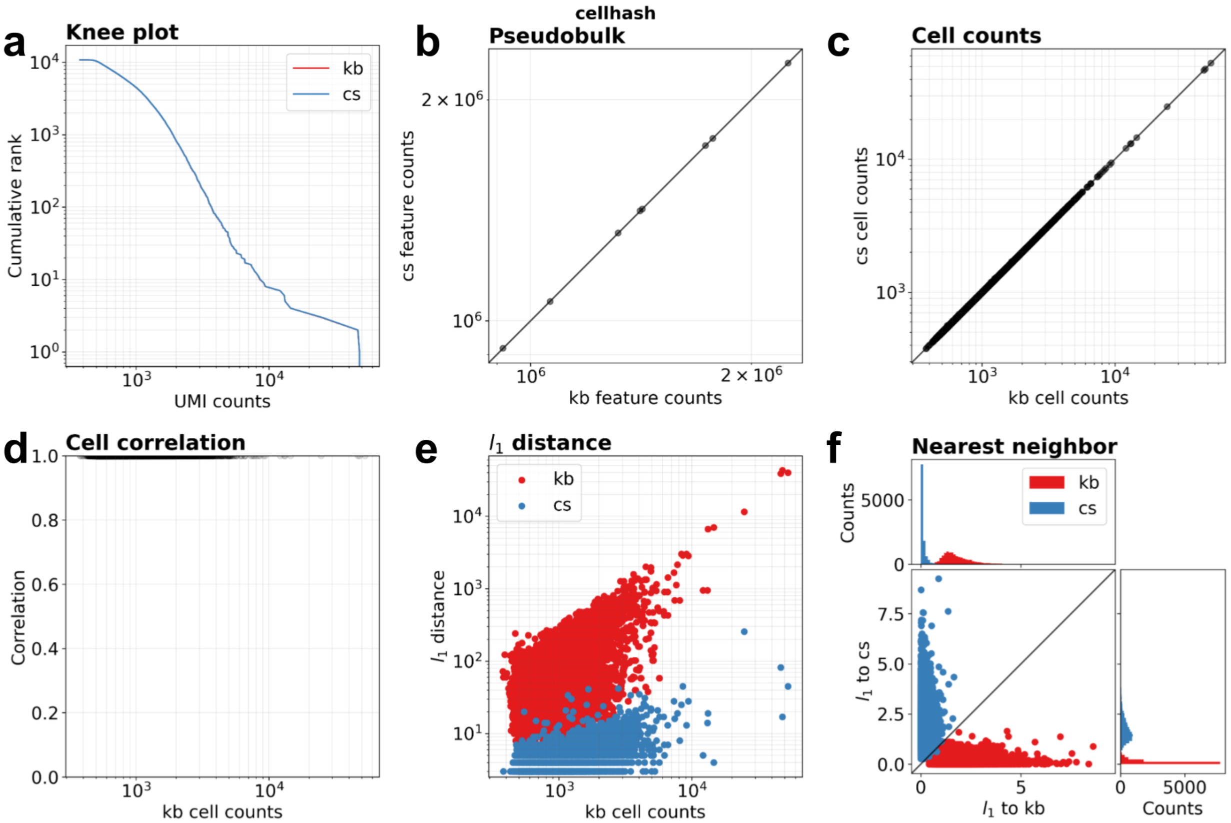


**Supplementary Figure 3:** Comparison between kallisto bustools (kb) quantifications and CITE-seq-Count (cs) quantifications for the 10x Feature Barcoding assay. **(a)** Knee plot comparing cumulative UMI counts per cell. **(b)** Pseudobulk comparison of cumulative UMI counts per feature barcode. **(c)** Cumulative UMI counts per cell. **(d)** Pearson correlation of the same cell between the two quantifications. **(e)** The l_1 distance between a kallisto bustools cell and its CITE-seq-Count doppelganger (blue) and the same cell and its nearest kallisto bustools neighbor (red) across the total UMI counts for that cell. **(f)** The l_1 distance of a kallisto bustools cell (red) to its CITE-seq-Count doppelganger (y-axis) and to its nearest neighbor (x-axis) and the l_1 distance of a CITE-seq-Count cell (blue) to its kallisto bustools doppelganger (x-axis) and its nearest neighbor (y-axis). The marginal distributions show that each kallisto bustools cell is closest to its corresponding CITE-seq-Count cell and that each CITE-seq-Count cell is nearest to its corresponding kallisto bustools cell.

a

b

```
# Multiseq
BC49: GACCAGCC
BC74:  ACCAGCCG

# 10xCRISPR
Protospacer: AAGCAGTGGTATCAACGCAGAGTACATGGGG -(BC)- GTTTAAGAGCTAAGCTGGAAACAGCATAGCAAGTTTAAAT

EZR-1:    CACTCGGCGGACGCAAGGG
EZR-2: GCGCACTCGGCGGACGCAA

PPIB-1: GGAGAGGCGCAGCATCCAC
PPIB-2:    GAGGCGCAGCATCCACAGG
```

**Supplementary Figure 4: (a)** Multiseq barcodes that share a 7 bp subsequence, **(b)** 10xCRISPR screen barcodes that share a 16 bp subsequence and share sequence similarity to the end of the first protospacer sequence fragment.

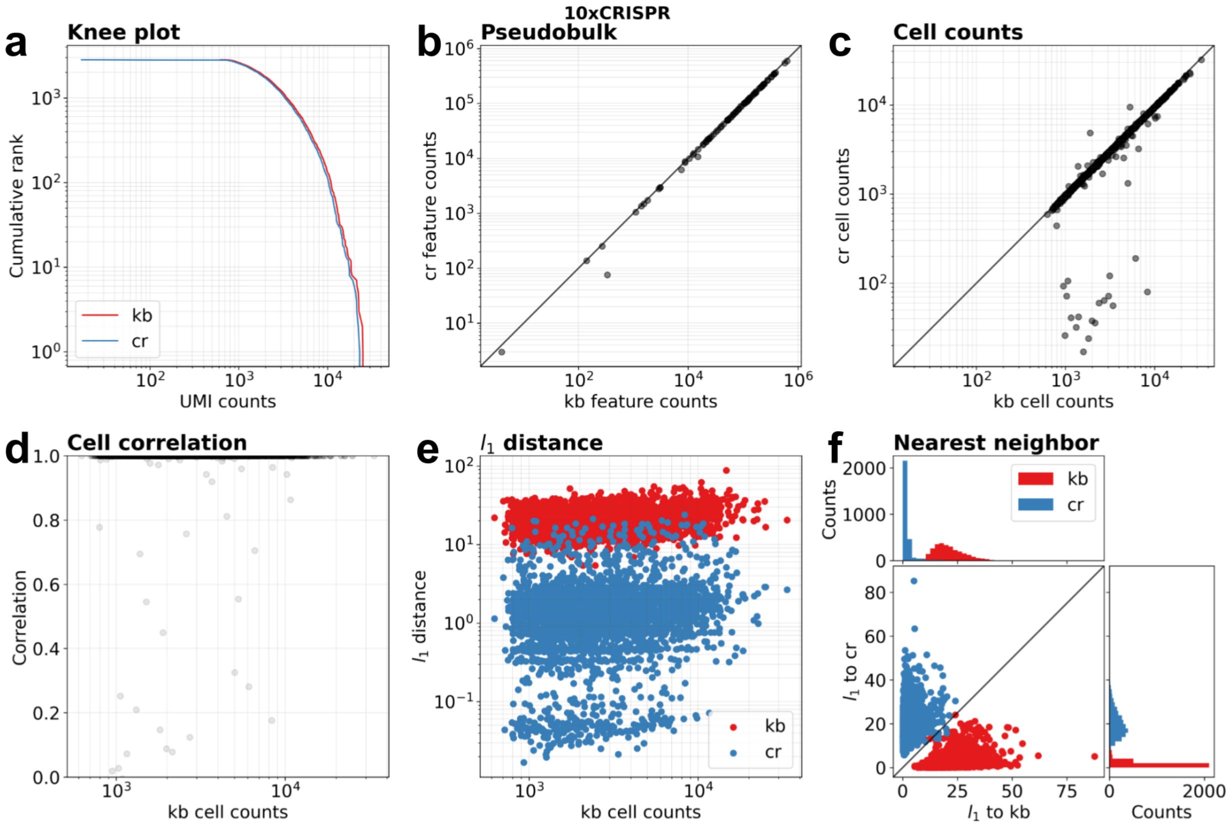


**Supplementary Figure 5:** Comparison between kallisto bustools (kb) quantifications and Cell Ranger (cr) quantifications for the 10x Feature Barcoding assay. **(a)** Knee plot comparing cumulative UMI counts per cell. **(b)** Pseudobulk comparison of cumulative UMI counts per feature barcode. **(c)** Cumulative UMI counts per cell. **(d)** Pearson correlation of the same cell between the two quantifications. **(e)** The l_1 distance between a kallisto bustools cell and its Cell Ranger doppelganger (blue) and the same cell and its nearest kallisto bustools neighbor (red) across the total UMI counts for that cell. **(f)** The l_1 distance of a kallisto bustools cell (red) to its Cell Ranger doppelganger (y-axis) and to its nearest neighbor (x-axis) and the l_1 distance of a Cell Ranger cell (blue) to its kallisto bustools doppelganger (x-axis) and its nearest neighbor (y-axis). The marginal distributions show that each kallisto bustools cell is closest to its corresponding Cell Ranger cell and that each Cell Ranger cell is nearest to its corresponding kallisto bustools cell.

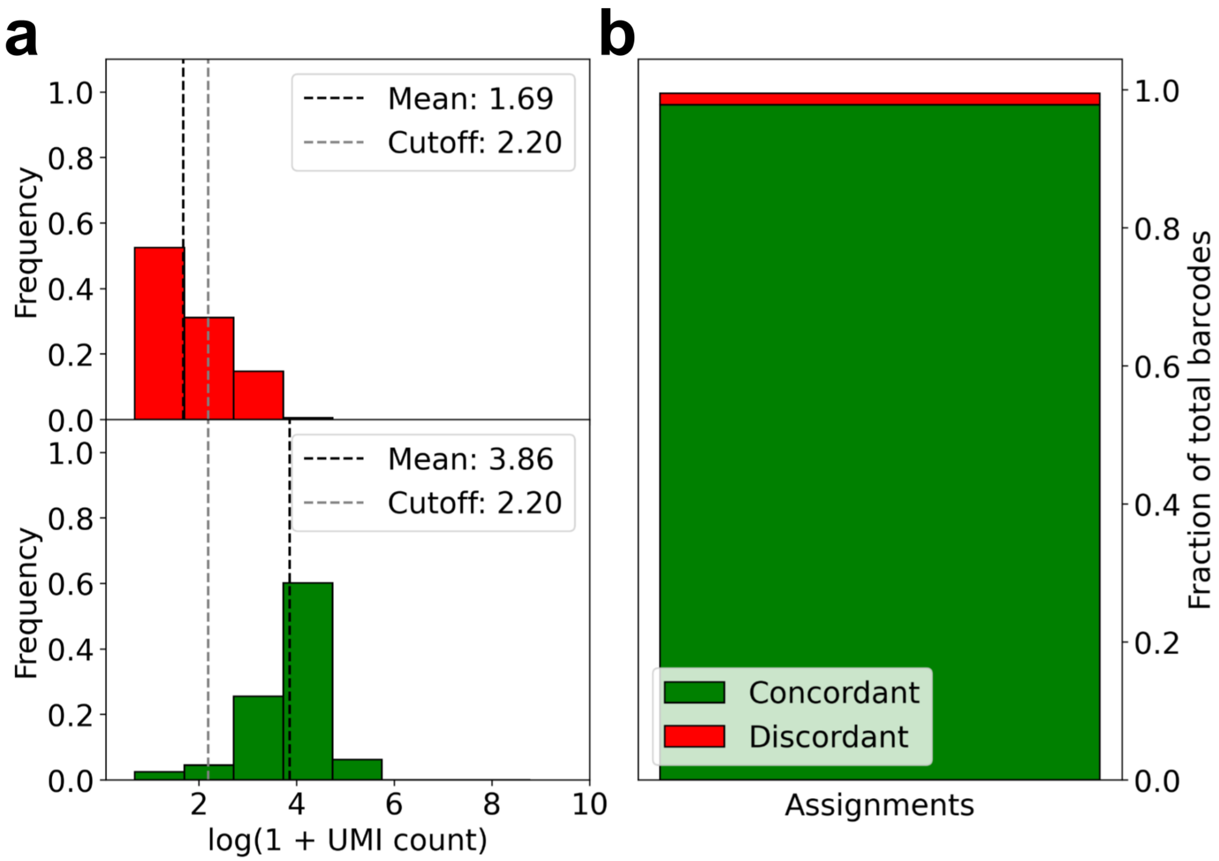


**Supplementary Figure 6: TAP-Seq barcode assignments. (a)** The UMI count distribution on cells that are fully discordant with the TAP-Seq assignments (red) and fully concordant with the TAPseq assignments (green). The mean log1p UMI count (black dotted line) and log1p UMI cutoff (gray dotted line) are plotted. **(b)** The fraction of total barcodes that are fully concordant and fully discordant with the TAP-Seq assignments

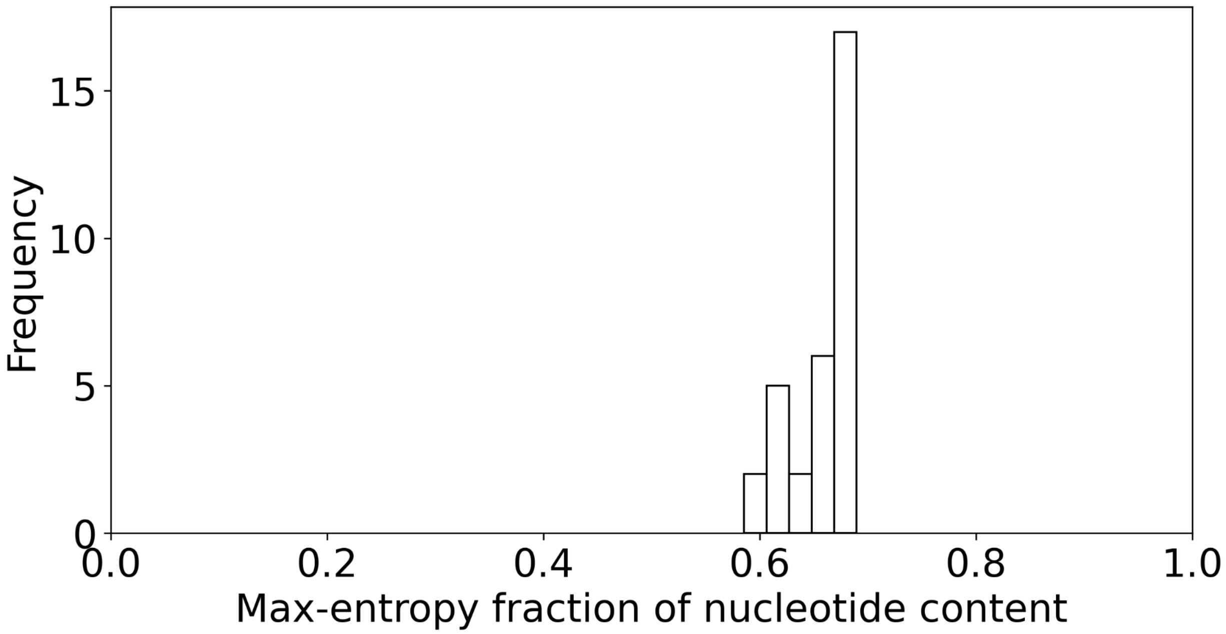


**Supplementary Figure 7:** The entropy of the per barcode distribution on base pair frequency with respect to the maximum entropy.

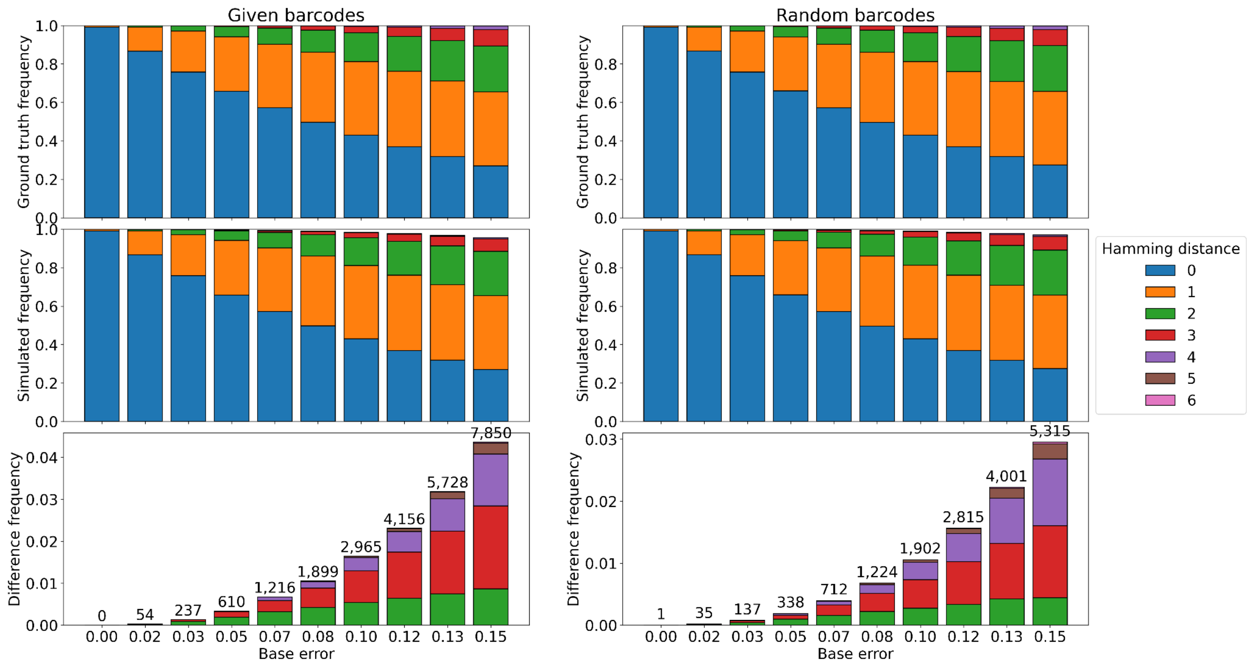


**Supplementary Figure 8: Barcode simulation**. For a given barcode from a set of barcodes we generate a collection of barcode mutants. We then check to see how many mutants that were originally hamming distance X apart are ambiguous, after creating errors in the bases. We perform this simulation for a supplied list of barcodes (left) and a random list of barcodes, the same length and the same number as those provided (right).

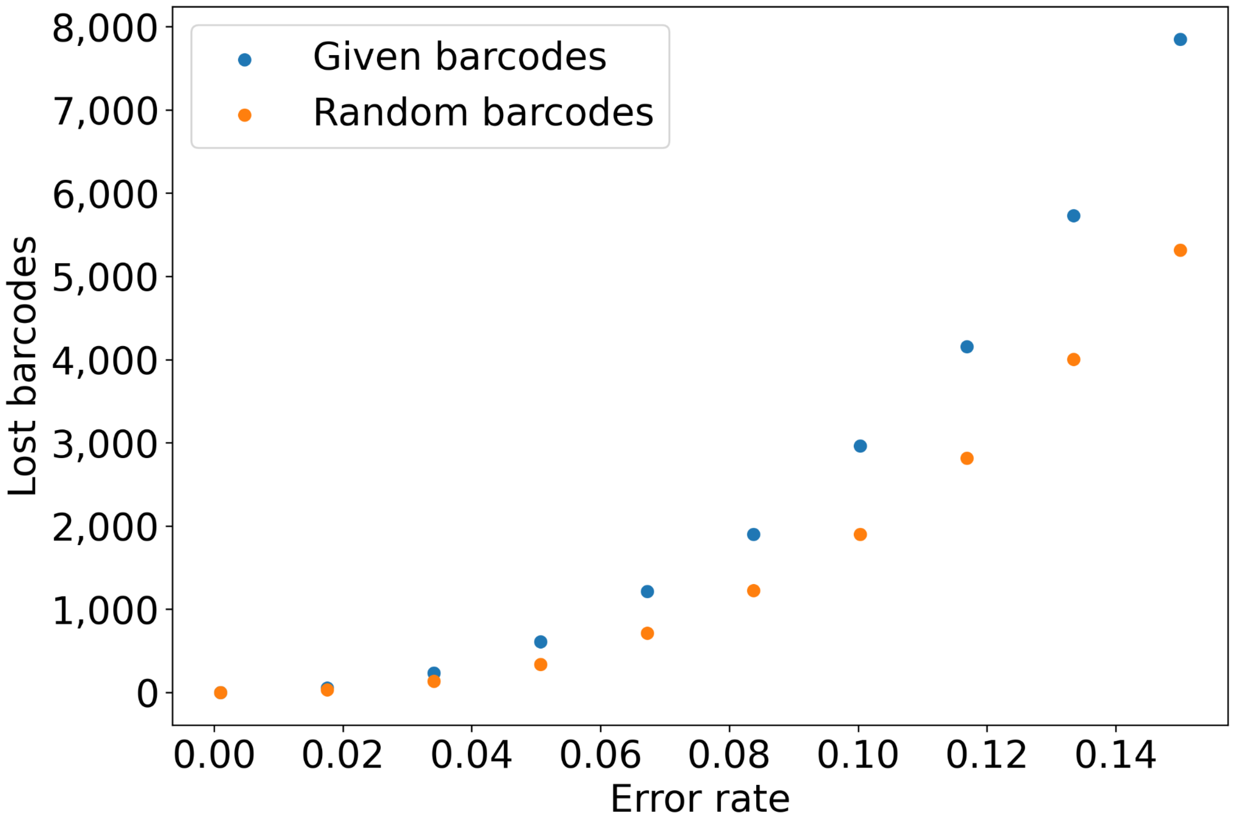


**Supplementary Figure 9: Barcode simulation of Multiseq and random barcodes.** Mutant barcodes are simulated against the Multiseq barcodes (blue) and random barcodes (orange) for a given per-base error rate. Mutant barcodes are error-corrected and the amount of "lost" barcodes that cannot be unambiguously corrected, to a given barcode of origin, are counted and compared.

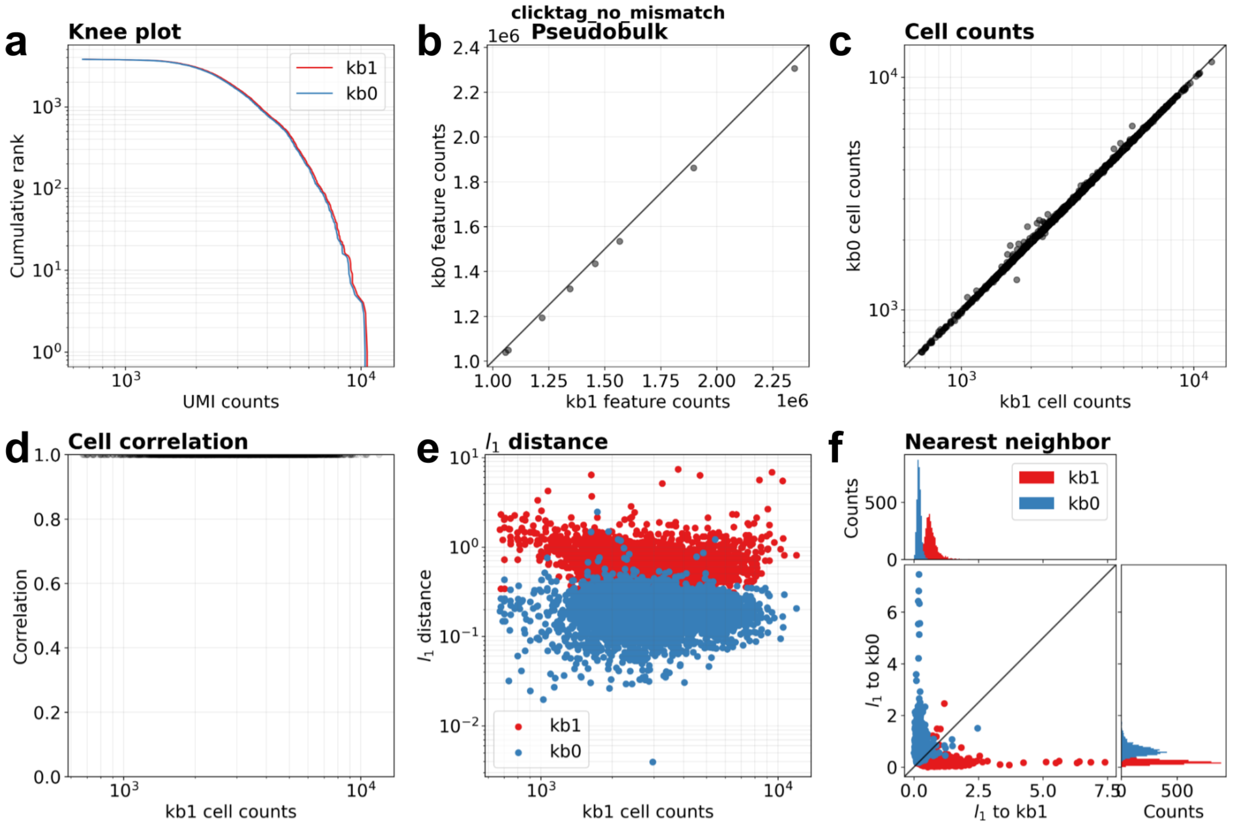


**Supplementary Figure 10:** Comparison between kallisto bustools quantifications with the hamming-1 mismatch index (kb1) and the kallisto bustools quantifications with the hamming-0 mismatch index (kb0) for the Clicktag assay. **(a)** Knee plot comparing cumulative UMI counts per cell. **(b)** Pseudobulk comparison of cumulative UMI counts per feature barcode. **(c)** Cumulative UMI counts per cell. **(d)** Pearson correlation of the same cell between the two quantifications. **(e)** The l_1 distance between a kallisto bustools-1 cell and its kallisto bustools-0 doppelganger (blue) and the same cell and its nearest kallisto bustools-1 neighbor (red) across the total UMI counts for that cell. **(f)** The l_1 distance of a kallisto bustools-1 cell (red) to its kallisto bustools-0 doppelganger (y-axis) and to its nearest neighbor (x-axis) and the l_1 distance of a kallisto bustools-0 cell (blue) to its kallisto bustools-1 doppelganger (x-axis) and its nearest neighbor (y-axis). The marginal distributions show that each kallisto bustools-1 cell is closest to its corresponding kallisto bustools-0 cell and that each kallisto bustools-0 cell is nearest to its corresponding kallisto bustools-1 cell.
